# Structural characteristics of important daily movement corridors for waterbirds in urban areas: a field test focusing on Black-headed Gulls

**DOI:** 10.64898/2026.09.01.748747

**Authors:** Shiori Takeshige, Yusuke Sawa, Kazuhiro Katoh

## Abstract

1. Many global cities have developed along rivers, subjecting riverine ecosystems to extensive channelization. Straightened rivers have typically been considered too modified to be suitable for aquatic organisms and thus have lacked conservation priority. But considering avian movement decision, such as distance savings and obstacle avoidance, straightened river channels may function as critical movement corridors. Consequently, re-evaluating the ecological value of channelized rivers based on movement ecology can benefit urban waterbird conservation.

2. We tracked the daily movement behaviors of the Black-headed Gull (*Chroicocephalus ridibundus*), a river-dependent waterbird distributed across the Palearctic, and identified key movement corridors in highly urbanized Tokyo, Japan. We collected GPS tracking data for three years from 35 individual gulls within three regions of central Tokyo and additionally measured variables that likely affect bird navigation: river sinuosity and urban feature height (i.e., buildings). We then used generalized linear mixed models to determine relationships between these variables with distances to river centerline (representing fidelity to the river during navigation) across various temporal scales reflecting different individual decision times. Additionally, we predicted and mapped these distances to river centerline across individual river channels in central Tokyo to identify important movement corridors.

3. River structure and feature heights influenced individual river fidelity, but this depended on the spatial and temporal scale of decision-making. Higher feature heights promoted river fidelity during movements based on sudden and mid-term decisions, but not long-term decisions. Conversely, under the broadest perceptual range and for long-term decisions, channel straightness significantly enhanced river fidelity. Overall, results showed that straight river channels with high feature heights represent crucial movement corridors for gulls.

4. Our model identified a key corridor in central Tokyo for waterbird movement that serves to connect daytime habitats to roosting sites. This predictive tendency was also highly robust across all time scales.

5. While straightened river channels with high feature heights may provide unsuitable habitats for some taxa, our results demonstrate their ecological value and importance for conservation as essential movement corridors for waterbirds.

## Introduction

As many cities globally have developed along rivers, riverine ecosystems have undergone severe anthropogenic modification (Forman, 2014; Chin, 2006). Among these modifications, channelization is a primary driver degrading rivers’ functionality as habitats. Straightened river channels homogenize flow velocities and eliminate variations in water depth and microhabitats, resulting in simplified environments (Rhoads et al., 2003). Consequently, suitable conditions for the establishment and persistence of diverse organisms, are drastically reduced (Rhoads et al., 2003; Nakano & Nakamura 2006, 2008; Nakamura et al., 2014). Conversely, meandering rivers exhibit high environmental heterogeneity, characterized by alternating riffle-pool structures and lateral depth gradients. Such dynamic structures generate diverse aquatic habitats, such as oxbow lakes and backwaters, which serve as indispensable fish habitats (Winemiller et al., 2000; Nagayama & Nakamura 2018). Therefore, traditional habitat conservation and restoration frameworks have heavily emphasized the ecological value of meandering rivers, prompting widespread channel remeandering projects worldwide (e.g. Moerke et al., 2004; Pedersen et al., 2007; Lorenz et al., 2009).

Recently, however, urban rivers have increasingly been recognized not only as foraging or breeding habitats, but also as vital movement corridors for wildlife. Studies focusing on urban waterbird movements demonstrate that rivers function as critical movement pathways (Skórka et al., 2009), a tendency that is especially pronounced in highly urbanized landscapes (Takeshige & Katoh 2025). Waterbirds, predominantly gulls and cormorants, fly along rivers (Takeshige & Katoh 2020), with higher abundances of moving individuals observed in rivers with larger open-water areas; conversely, artificial structures covering rivers can impede waterbird flight and restrict corridor functionality (Takeshige & Katoh 2023). Furthermore, research investigating how surrounding landscape urbanization influences corridor function reveals that Black-headed Gulls, which rely heavily on rivers for movement, tend to fly along rivers more frequently in more urbanized areas (Takeshige et al., 2025). Because moving waterbirds facilitate essential ecosystem services—such as seed dispersal, transport of small organisms, and nutrient loading—securing their movement pathways is vital for sustaining urban ecosystem functioning (Green & Elmberg 2014). Despite the recognized importance of rivers as waterbird movement spaces, the specific characteristics that render particular river channels crucial for their movement remain unknown.

To identify river channels crucial for movement, research on terrestrial animal movements in fragmented landscapes offers valuable insights. Animals frequently take shortcuts across the surrounding matrix when doing so substantially reduces their overall travel distance (St. Clair et al., 1998; Bélisle & Desrochers 2002). In riverine systems, navigating along a sinuous channel increases travel distance compared to a straight segment when moving between two points separated by the same straight-line distance. Consequently, following sinuous channels imposes higher energetic costs due to the extended travel distance. In addition to channel sinuosity, the height of surrounding landscape features is expected to dictate a river’s importance as a movement corridor. This is because ascending flight to clear obstacles via flapping consumes substantially more energy in birds than steady horizontal flight (Berg & Biewener 2008). Therefore, when feature heights inside a river bend are low, birds can achieve distance savings while maintaining horizontal flight by cutting across the terrestrial matrix inside the bend. Under such conditions—where channel sinuosity is high and interior feature heights are low—the functional importance of the river channel as a movement corridor likely diminishes. Taken together, we expect that channel sinuosity (which governs horizontal travel distance) and surrounding feature heights (which dictate vertical flight adjustments) jointly define the quality of river channels as movement corridors. Although movement corridor research has predominantly focused on terrestrial organisms (e.g. Gillies et al., 2011; Tremblay & St. Clair 2011) and informed practical urban planning (e.g., European Commission, 2022), evaluating corridor importance based on such avian energetic movement decision has remained unexplored.

This study elucidates the characteristics of critical river channels based on movement decision of waterbirds. To this end, we tracked movement behavior of Black-headed Gulls (*Chroicocephalus ridibundus*), a globally distributed species known to rely heavily on rivers as movement routes, in densely populated Tokyo, Japan. Black-headed gulls are common wintering waterbirds in Tokyo that forage along rivers and nearby ponds and roost in Tokyo Bay. Therefore, maintaining connectivity between inland waters and the coast is crucial for them. We elucidated the local conditions of river channels where gulls flew. We then used these results to map potential key river channels for waterbird movement in this hyper-urban environment. Ultimately, this research aims to provide valuable insights for waterbird conservation in highly urbanized landscapes.

## Materials and Methods

### Study area

The study area is located in Japan, at the center of the metropolitan area along inner Tokyo Bay (Fig. 1). Several mid-sized urban rivers, including the Sumida, Arakawa, Nakagawa, Kyu-Edogawa, Edogawa, and Tamagawa rivers, flow through the heart of the Tokyo metropolitan area, with urban land-use expanding across their watersheds. Along these river basins, at least 17 ponds serve as wintering sites for waterbirds (Takeshige & Katoh 2023). Furthermore, Kasai Rinkai Park, a Ramsar site, is located at the estuaries of the Arakawa and Kyu-Edogawa rivers, while Sanbanze—a tidal flat that serves as a critical stopover and wintering site for migratory waterbirds—is situated at the mouth of the Edogawa River.

**Fig 1.**
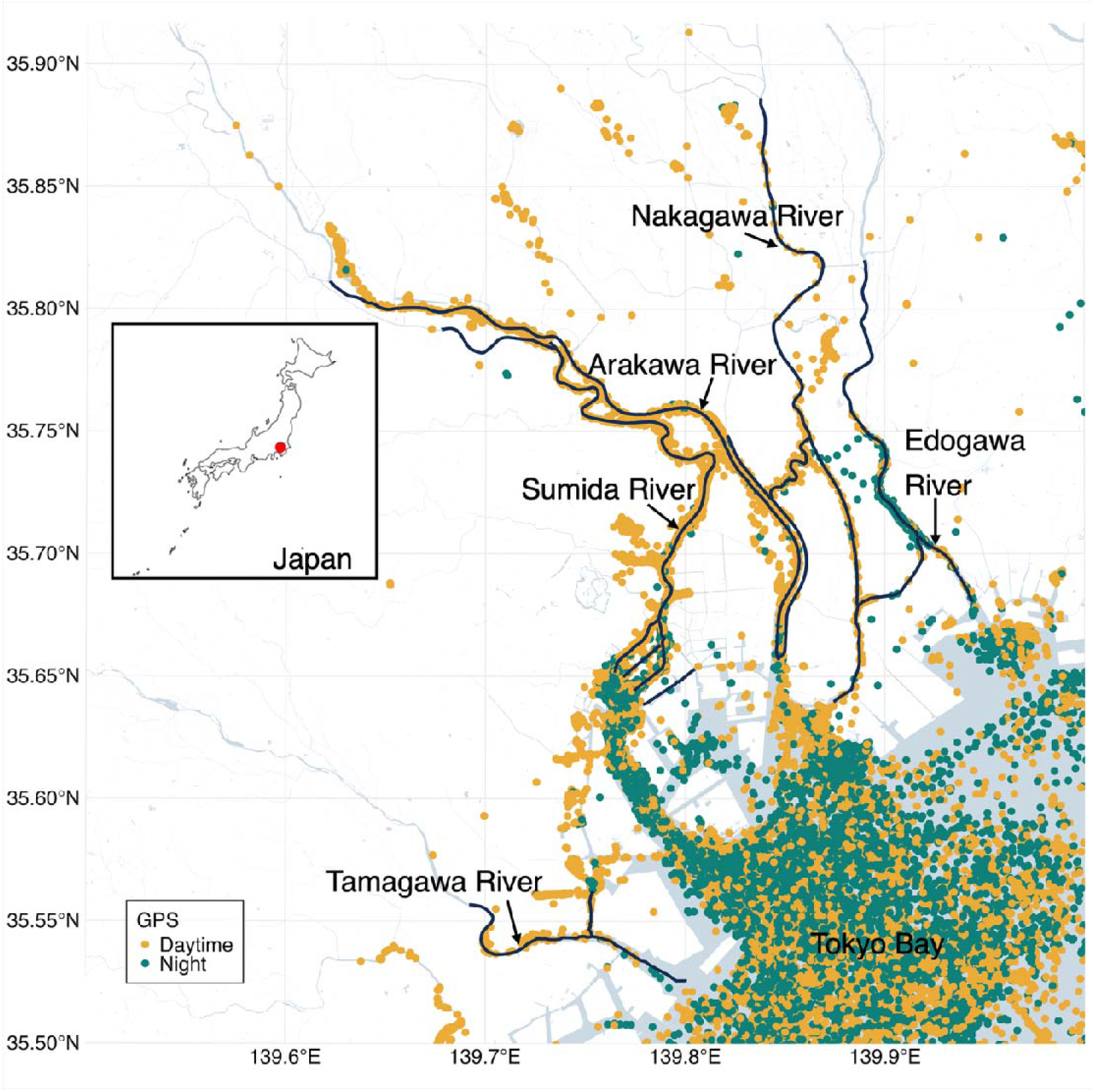
Map showing the study area and GPS data from Black-headed Gulls. Black lines represent the targeted river segments for which the CD (closest distance to the river) was estimated. Among the rivers in central Tokyo, those within the spatial coverage of the moving gull GPS location data were selected. Data collected between 05:00 and 16:59 were defined as daytime, whereas data from 17:00 to 04:59 (all remaining hours) were classified as night. We show only one capture site (Sumida River) because as a result of data processing steps, data from only 13 individuals captured in the Sumida River were extracted.

### GPS relocation data

Observational studies have demonstrated that *C. ridibundus* exhibits a high dependence on rivers as movement pathways (Takeshige & Katoh 2020; Takeshige & Katoh 2023), and this tendency becomes stronger in more highly urbanized areas (Takeshige et al., 2025). We captured 35 gulls from three areas: Sumida River (n = 23), Teganuma Lake (n = 4), and Senbako Lake n = 8) from 2021 to 2023. Detailed information regarding the captured individuals is provided in Takeshige et al. (2025). We attracted individuals with bread and captured them carefully by hand. All birds were fitted with metallic color rings (0.76 g and 0.7 g, respectively) and a GPS logger (Druid Technology, Debut NANO, 3.7 or 3.8 g, China) attached to a harness. For individuals whose body mass was recorded, the total weight of attachments was less than 3% of their body mass, a threshold known to have no effect on seabird behavior (Phillips et al., 2003). The standard data collection interval for the GPS loggers was set to 1 h for each individual. In addition, we configured the following high-resolution intervals to save battery life: (1) 600 s when voltage exceeded 3.97 V, (2) 120 s when it exceeded 4.02 V, and (3) 60 s when it exceeded 4.10 V. Furthermore, a 60-s interval was triggered if flight speed exceeded 5 m/s while the voltage was above 3.92 V; this function ceased when the speed decreased below 2 m/s. To ensure the accuracy of the location data, we restricted the dataset to GPS fixes that were recorded by three or more satellites. All data subjected to this preprocessing step are summarized in Fig. 1. The obtained location fixes were converted into a series of regular steps (hereafter referred to as a “burst”) using the R package *amt* (Signer et al., 2019). Through this process, the sampling interval of the GPS data was standardized to 60 s with a tolerance of 30 s. A burst was defined as a sequence of consecutive locations at this sampling interval, while a step was defined as the displacement between two consecutive locations. Bird capture and tracking procedures were approved by the Ethics Committee of our institution and conducted under scientific permits granted by governments located in capture sites.

Because our primary interest focused on movements near rivers, we first excluded steps from each burst that fell below the lower 5% threshold of the gamma distribution of step lengths; this was done to eliminate localized, circular movements within the same area. The gamma distribution is commonly used to model the distribution of step lengths. Next, we removed bursts that did not contain at least two steps within the study area. Furthermore, to filter out initial soaring behavior upon departure and settling behavior upon arrival at destinations, we excluded GPS locations within 500 m of the start point until the track moved beyond this radius, as well as locations within 500 m of the end point of each burst. We also removed GPS locations that were recorded within lakes and ponds. Finally, bursts that were reduced to two or fewer steps after these screening procedures were excluded.

### Spatial covariates

To identify river channels crucial for movement, we extracted river sinuosity and the height of features along the river as characteristics of each segment. In this paper, a “river segment” refers to each of the equal-length sections into which the river is divided, from the river mouth up to an arbitrary extent, along the river centerlines. The sinuosity of a river segment directly affects the travel distance to a destination. If following the river results in an excessive detour (i.e., when the segment is highly sinuous), animals may choose to take a shortcut. Additionally, the height of features along the river influences the frequency of vertical flight adjustments. When the feature height along a river is high, deviating from the river requires a steep ascent, which increases energy expenditure. We established three spatial scales (1, 2, and 5 km) to represent the river segments perceived or remembered by the gulls. Because the exact scale at which gulls make decisions remains unclear, we evaluated three distinct sizes. For each segment size, we calculated river sinuosity and the height of features along the river (the mean value of the Digital Surface Model, DSM). River sinuosity was defined as the channel length along the river centerline from the start to the end of the segment, divided by the straight-line distance between the same start and end points. To calculate the height of features along the river, a 500-meter buffer was generated on both sides of the river centerline, and the mean DSM value within this buffer was calculated. Any negative DSM values were treated as missing data due to data accuracy concerns. Water area polygon data from the Geospatial Information Authority of Japan (2022) was used to calculate river sinuosity. For the DSM data, we utilized the ALOS World 3D – 30 m dataset (Japan Aerospace Exploration Agency, 2025). We performed processing and manipulation of data as well as all spatial and statistical analyses using R, version 4.4.2 (R Core Team, 2026).

### Data analysis

The overall analytical workflow of this study is illustrated in Fig. 2. To identify key river segments for movement while accounting for gull movement decision, we evaluated how river sinuosity and the height of features along the river influence the extent to which movement trajectories follow the river. We divided bursts into movement segments of 1 step, and consecutive windows of 2 steps, 3 steps, 5 steps, and 10 steps. Here, a “movement segment” is defined as a sequence within a burst that reflects the temporal scale at which individual gulls make movement route decisions. The five movement segment scales were established to capture both instantaneous and mid- to long-term decision-making processes. At each movement segment scale, we analyzed the effects of river segment characteristics on river use as a movement pathway. The closest distance (CD) to the river centerline, representing the extent to which movement segments follow the river, was calculated using Equation 1:

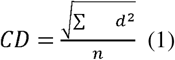

where *d* is the distance from the river centerline to each location point within the movement segment, and *n* is the number of location points. Higher CD values indicate that the individual gull flew further away from the river, or in other words, it had lower “river fidelity”.

**Fig. 2.**
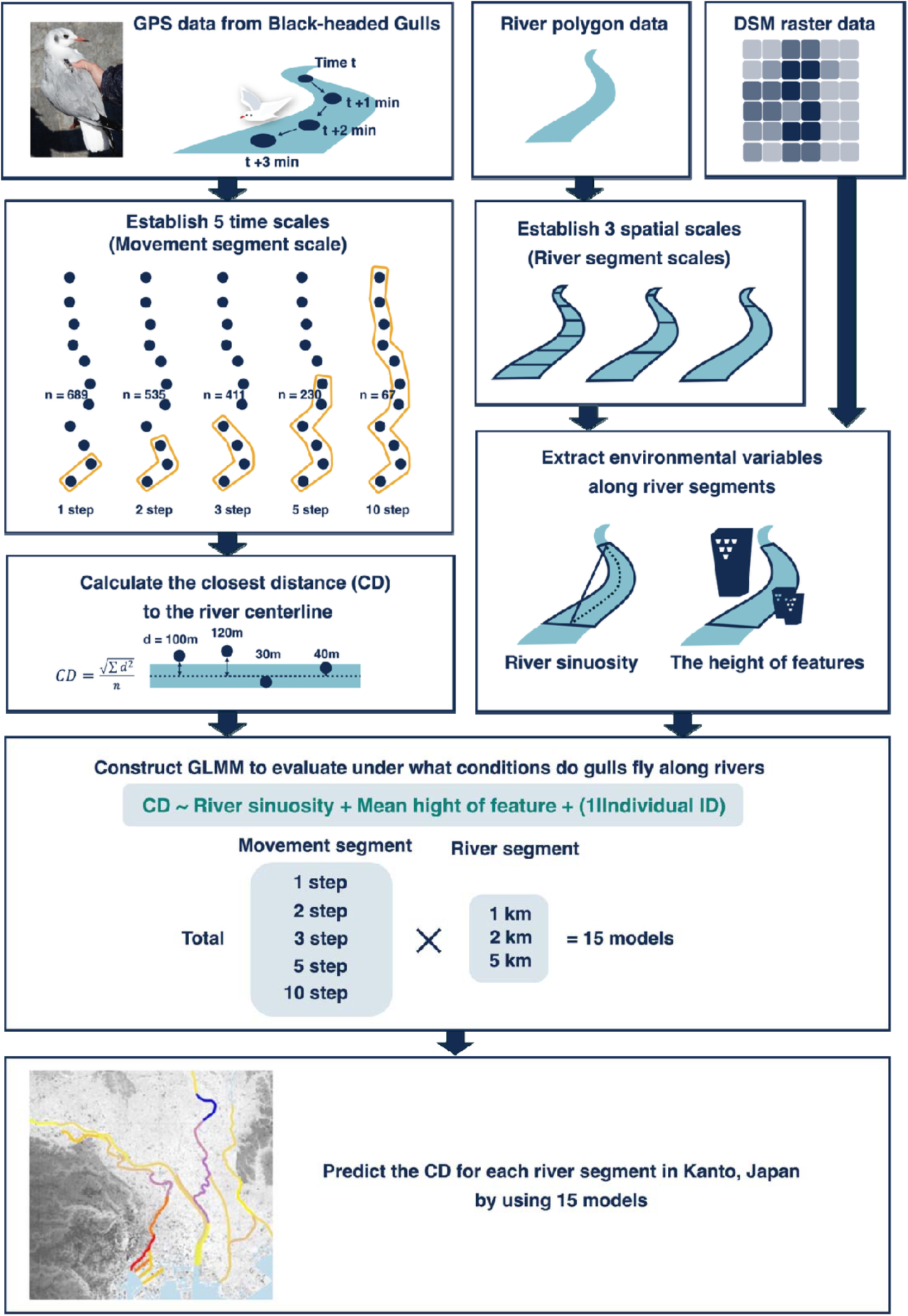
Conceptual illustration of the study design. GPS relocation data from Black-headed Gulls were divided into movement segments across five temporal scales (1, 2, 3, 5, and 10 steps), and closest distance (CD) values were calculated to quantify the extent to which movement segments follow the river. Here, d represents the distance from the river centerline to each location point within a movement segment, and n is the total number of location points. River characteristics (sinuosity and mean feature height calculated based on the Digital Surface Model, DSM) were evaluated across three spatial scales (1, 2, and 5 km) reflecting the gulls’ potential perceptual or memory ranges. Generalized Linear Mixed Models (GLMMs) were constructed for all combinations of movement segment scales and spatial river segment sizes to evaluate under what conditions gulls fly along rivers. Finally, predicted CD values were mapped across river segments in central Tokyo to identify critical movement corridors.

Using R version 4.4.2 (R Core Team, 2026), we constructed Generalized Linear Mixed Models (GLMMs) with the CD of each movement segment as the response variable, and both the sinuosity of the river segment containing the segment’s starting point and the mean height of features along that river segment as explanatory variables. Individual ID was included in the models as a random effect. These models were developed for all possible combinations of river segment size (1 km, 2 km, and 5 km) and movement segment scales (1 step, 2 steps, 3 steps, 5 steps, and 10 steps). All models were fitted using the glmer function in the lme4 package version 1.1.38 (Bates et al., 2015) and we performed likelihood ratio tests for the coefficients using the Anova function in the car package version 3.1.3 (Fox and Weisberg 2019). Furthermore, using these models, we predicted the CD for each river segment as an index of river fidelity (the extent to which movement segments follow the river). Among the river segments in central Tokyo, those within the spatial coverage of the gull GPS location data were selected.

## Results

We obtained a total of 214,500 GPS location fixes from 35 gulls. While gulls utilized inland waterbodies and coastal areas during the day, they primarily roosted at sea at night, and major rivers within the study area connected daytime habitats directly with nocturnal marine roosts as single continuous waterbody (Fig. 1). Our sequence of filtering processes extracted 94 movement bursts, comprising 875 movement locations from 13 individuals captured in the Sumida River. From these data, we extracted movement segments corresponding to 1 step (n = 689) as well as consecutive windows of 2 (n = 535), 3 (n = 411), 5 (n = 230), and 10 steps (n = 67).

Individual fidelity to the river centerline was influenced by both river sinuosity and the height of features along rivers. However, these effects varied depending on the timescale of movement (Fig. 3, Table 1). Feature height had a significantly negative correlation with CD for all spatial scales (1, 2, and 5 km) and nearly all temporal movement scales (1, 2, 3, and 5 steps), with the exception of the 10-step range, for which all spatial scales were not significant. On the other hand, river sinuosity had a significantly positive relationship at 5 km for all temporal movement scales, with the addition of 2 km for the long-term 10-step range. As the temporal movement scale increased from 1 to 5 steps, the coefficient values of feature height decreased (Fig. 3). Conversely, the coefficient values of river sinuosity at the long-term scale (10 steps) were approximately twice as large as the short- to mid-term scales (e.g., from 0.07 at 1-step to 0.14 at 10-step at 5 km scale). The random intercept for individual identity accounted for minimal variance in the models, with estimates ranging from 0.06 to 0.25 relative to the fixed intercepts of 4.30 to 4.56.

**Fig. 3.**
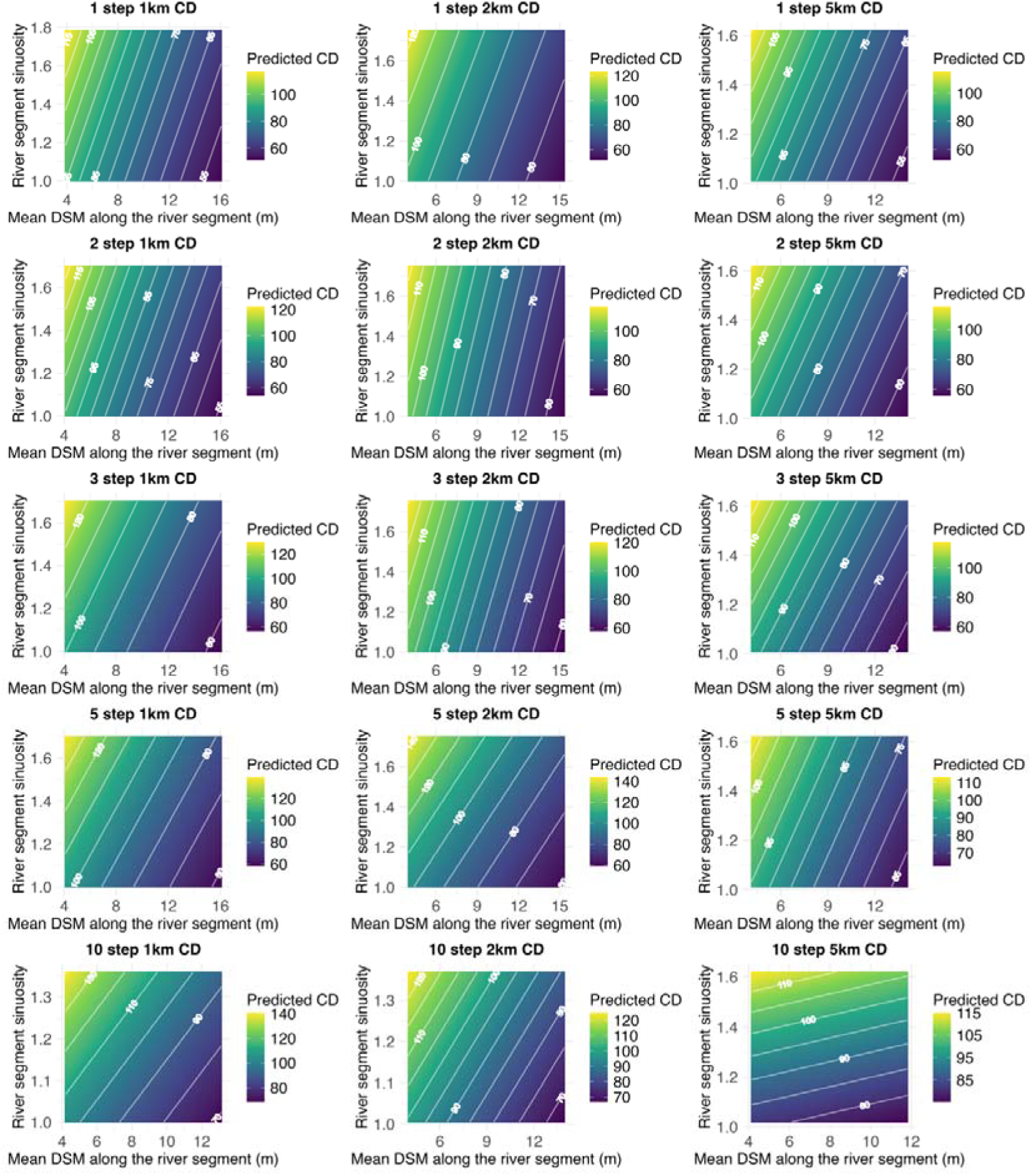
Closest distance to the river (CD) values at different movement segment scales and river segment sizes. CD value represents the extent to which gull’s movement segments follow the river. DSM (Digital Surface Model) represents the height of features along the river. Higher CD values indicate that gulls fly further away from the river they are navigating. The color gradient represents the predicted CD values based on the GLMM for each movement segment scale and river segment size. This result indicates that gulls tended to fly along rivers within straight segments characterized by high feature heights.

**Table 1.** Coefficients of GLMM and the results of likelihood ratio tests constructed for each movement segment and river segment. These models were developed for all possible combinations of river segment size (1 km, 2 km, and 5 km) and movement segment scales (1 step, 2 steps, 3 steps, 5 steps, and 10 steps). River segment refers to each of the equal-length sections into which the river is divided, from the river mouth up to an arbitrary extent and represents spatial scales perceived or remembered by the gulls. Movement segment is defined as a sequence within a series of regular steps that reflects the temporal scale at which individual gulls make movement route decisions.

| Model |  | Intercept | Random effect<br>Std.Dev | Mean feature<br>height along<br>river segment | (p value) | Sinuosity of<br>river segment<br>(with starting<br>point) | (p value) |
| --- | --- | --- | --- | --- | --- | --- | --- |
| 1 step | 1km | 4.3 | 0.13 | -0.19 | p<0.05 | 0.02 | 0.36 |
|  | 2km | 4.31 | 0.13 | -0.21 | p<0.05 | 0.03 | 0.21 |
|  | 5km | 4.31 | 0.14 | -0.17 | p<0.05 | 0.07 | p<0.05 |
| 2 step | 1km | 4.36 | 0.1 | -0.19 | p<0.05 | 0.02 | 0.32 |
|  | 2km | 4.36 | 0.1 | -0.19 | p<0.05 | 0.02 | 0.41 |
|  | 5km | 4.36 | 0.1 | -0.15 | p<0.05 | 0.07 | p<0.05 |
| 3 step | 1km | 4.4 | 0.06 | -0.17 | p<0.05 | 0.03 | 0.23 |
|  | 2km | 4.4 | 0.07 | -0.18 | p<0.05 | 0.02 | 0.33 |
|  | 5km | 4.4 | 0.07 | -0.14 | p<0.05 | 0.08 | p<0.05 |
| 5 step | 1km | 4.47 | 0.15 | -0.16 | p<0.05 | 0.04 | 0.23 |
|  | 2km | 4.47 | 0.16 | -0.15 | p<0.05 | 0.06 | 0.06 |
|  | 5km | 4.46 | 0.17 | -0.11 | p<0.05 | 0.06 | p<0.05 |
| 10 step | 1km | 4.56 | 0.24 | -0.08 | 0.07 | 0.06 | 0.13 |
|  | 2km | 4.56 | 0.23 | -0.07 | 0.11 | 0.09 | p<0.05 |
|  | 5km | 4.52 | 0.25 | -0.02 | 0.59 | 0.14 | p<0.05 |

Based on the CD model, we estimated the extent of river fidelity within each river segment and found that CD values varied substantially even among river corridors connecting daytime habitats and coastal roosts (Fig.4). Within straight segments with high DSM values (such as the lower reaches of the Sumida River that connect daytime habitats with roost habitats, Fig 1), gulls tend to fly along rivers, regardless of the movement segment scale or the river segment size (Fig. S1). At the 5-km river segment scale, models across all temporal scales predicted that the lower reach of the Sumida River had the lowest CD values, which were approximately half (0.47 to 0.66) of those recorded in the highest-value segments. Conversely, within the upper reaches of the Sumida River and the other rivers, gulls fly farther away from rivers compared with the lower Sumida River in almost all cases (Fig.4, Fig. S1).

**Fig. 4.**
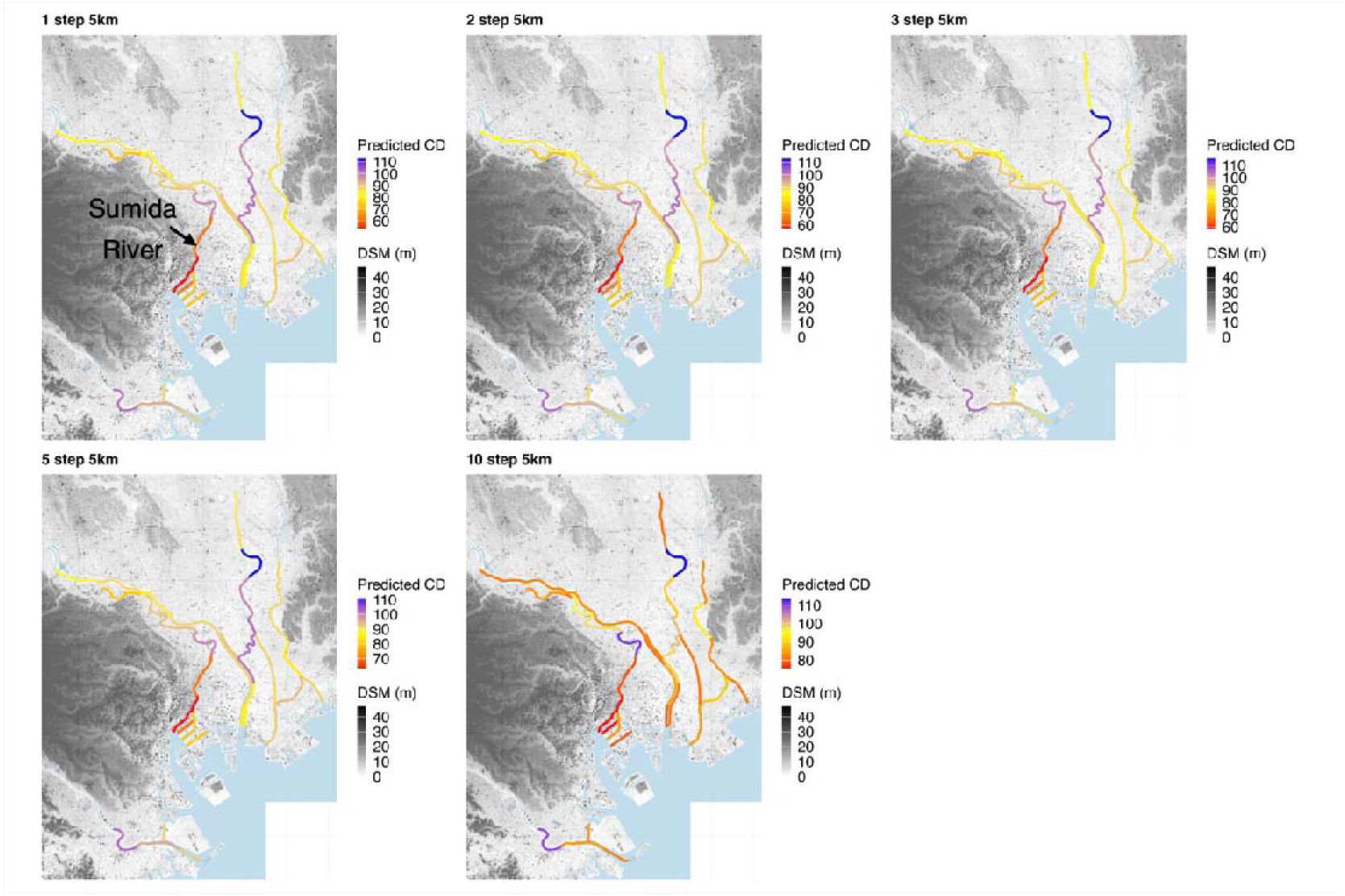
Predicted closest distance to the river (CD) values at each river segment. Results are shown for the 5-km river segment size, at which both the sinuosity and the mean DSM along the river segments were significant in most models. Warmer colors indicate a higher likelihood of gulls flying along the river segments. Regardless of the time scale of decision-making, river segments where gulls were more likely to fly along the river remained similar.

## Discussion

To our knowledge, this is the first study to demonstrate that straightened river channels—traditionally assumed to offer negligible contributions to biodiversity—can play a pivotal role as functional movement corridors for wildlife. In riverine landscapes, habitat heterogeneity is widely recognized as a paramount driver of biodiversity (Ward, 1998). Consequently, channelized or straightened river channels generally exhibit lower environmental heterogeneity, which in turn diminishes species richness (Nakano & Nakamura 2006, 2008; Nakamura et al., 2014). Indeed, environmental heterogeneity is frequently highlighted as a vital habitat characteristic for waterbirds (Andrade et al., 2018), and for this reason straightened river channels have seldom been prioritized in avian and broader biodiversity conservation strategies. In contrast, our findings reveal that straightened river channels are crucial for the daily movements of gulls. This outcome provides a novel perspective, showing that straightened river channels function effectively as connectivity pathways despite their limited capacity to support high biodiversity. Given that most urban gull movements are documented along rivers bridging daytime foraging sites and roosts (Fig. 1), conserving these river channels is crucial. These insights suggest that the ecological roles of urban rivers in supporting wildlife are heavily dictated by their spatial configurations, specifically channel sinuosity. Therefore, we propose that urban river management frameworks for waterbird conservation adopt a spatially differentiated strategy: heterogeneous, sinuous segments should be managed with an emphasis on habitat utilization, whereas the intervening straightened segments—or those bridging critical isolated waterbodies and alternative aquatic networks—should be maintained as movement corridors by preventing the installation of structures that impede waterbird flight pathways.

As expected, the importance of rivers as movement corridors was profoundly influenced by movement decision. With the increasing feasibility of tracking diverse species via devices such as GPS loggers, these findings strongly suggest that animal movement decision should be explicitly integrated into the identification of critical movement corridors. While the extent to which gulls fly along rivers was determined by river sinuosity and matrix characteristics, the magnitude of these effects varied across spatial and temporal scales. Gulls tended to avoid areas with high feature heights during instantaneous to mid-term decision-making in their daily movements, whereas they utilized straight river channels as movement routes for long-term decisions. Because an animal’s perceptual range during movement is rarely self-evident, as demonstrated in this study, we propose that corridor importance should be evaluated across diverse spatial and temporal scales to identify comprehensively crucial corridors.

Although straight river channels surrounded by relatively high features have been identified as important movement corridors, it is also essential to interpret these findings from the perspective of animal movement decision. There are two potential explanations for why gulls tended to fly along river channels in areas with higher feature heights. First, gulls may have been conserving energy by avoiding the costs associated with ascending over buildings. To clear these structures, gulls must climb using flapping flight, which demands a substantial amount of energy. Indeed, previous studies have demonstrated that ascending flight via flapping requires considerably more power than horizontal flight (Berg & Biewener 2008). Second, gulls may have been avoiding the atmospheric turbulence generated by buildings. Buildings are a primary driver of wind turbulence and the size increases with the scale of the structure (Shepard, 2025). Turbulence has been shown to increase the energetic costs of movement in many bird species, and it is widely suggested that navigating turbulent air imposes a heavy energetic burden, particularly on flapping fliers (Shepard, 2025). Given that areas with high feature heights in our study area are predominantly composed of tall buildings, gulls likely utilize river corridors to circumvent these aerodynamic challenges. One potential explanation for why gulls tended to fly away more from the river in highly sinuous segments is that they were conserving energy by shortening their travel distance. When flying between two points along a river, a more sinuous segment allows for greater distance savings if the animal cuts across the terrestrial landscape rather than following the river. Studies on terrestrial birds have demonstrated that birds are willing to cross a high-cost matrix if doing so allows them to substantially reduce their total travel distance (St. Clair et al., 1998; Bélisle & Desrochers 2002). In other words, the shape of the corridor dictates its usage, and river sinuosity serves as a crucial factor determining the importance of the river as a movement corridor.

Our findings offer valuable insights for conserving urban rivers as movement corridors for waterbirds. We suggest that straightened river channels with high feature heights could be considered for conservation as waterbird movement corridors; for example, the lower reaches of the Sumida River in Tokyo serve as an important movement corridor. Takeshige and Katoh (2023) showed that elevated roads running directly above rivers may impede bird flight along rivers, so to conserve these river channels, structures should be avoided that cover rivers in these segments or that obstruct bird movement, such as wind turbines and power lines. As urbanization expands in many areas of the globe, building heights along rivers are likely to increase. Under this scenario, even highly sinuous segments will likely become essential spaces for waterbird movement. A limitation of our study is that it focuses on only a single waterbird species, which stems from the fact that the species available for capture and tracking were highly constrained. As tracking and capture technologies advance in the future, it will become feasible to study a broader range of waterbird species. This will allow future research to clarify whether key river channels used as movement corridors vary depending on species traits and life histories.

## Acknowledgments

The authors thank Masumi Akimoto, Hitomi Abe, Yusaku Ohkubo, Yuta Okamoto, Noriyoshi Kawasaki, Kazuhiko Hirata, Emiko Sato, Nozomu J. Sato, Hideo Sato, Yusuke Takahashi, and Machi Hanawa for their help with the bird surveys. We also thank Hiromichi Suzuki for his insightful comments and fruitful discussions. We are also grateful to the Yamashina Institute for Ornithology for allowing us to use a sample of Black-headed Gulls to study how to attach GPS loggers. The authors also would like to thank Jamie M. Kass for his careful proofreading of the manuscript and for providing many helpful comments. This study was supported by a Sasakawa Scientific Research Grant 2020 from the Japan Science Society Grant Number 2020–5026, the Japan Society for the Promotion of Science KAKENHI Grant Number 22KJ2638, and partly by the Japan Society for the Promotion of Science KAKENHI Grant Number 17K08186 and Japan Science and Technology Agency Co-creation Place Formation Support Program Grant Number J240000159. We used DeepL and Gemini to proofread the manuscript text and analysis code, and we extensively reviewed and verified all outputs.

## Author Contributions

All authors conceived the ideas and designed the methodology; TS and SY collected the data; TS analyzed the data; and TS wrote the manuscript. All authors contributed critically to drafts and approved the final manuscript for publication.

## Data Availability

GPS relocation data from this study have been deposited in Movebank (Movebank ID: 9339187678). All data that support the findings of this study are available from the corresponding author upon reasonable request.

## References

Andrade, R., Bateman, HL., Franklin, J., & Allen D. (2018). Waterbird community composition, abundance, and diversity along an urban gradient. Landscape and Urban Planning, 170, 103–111. 10.1016/j.landurbplan.2017.11.003

Bates, D., M, Mächler., B, Bolker., & S, Walker. (2015). Fitting Lin-ear Mixed-Effects Models Using lme4. Journal of Statistical Software, 67(1), 1–48. 10.18637/jss.v067.i01

Bélisle, M., & Desrochers, A. (2002). Gap-crossing decisions by forest birds: an empirical basis for parameterizing spatially-explicit, individual-based models. Landscape Ecology, 17, 219–231. 10.1023/A:1020260326889

Berg, AM., & Biewener, AA. (2008). Kinematics and power requirements of ascending and descending flight in the pigeon (*Columba livia*). Journal of Experimental Biology, 211 (7), 1120–1130. 10.1242/jeb.010413

Chin, A. (2006). Urban transformation of river landscapes in a global context. Geomorphology, 79 (3-4), 460–487. 10.1016/j.geomorph.2006.06.033

European Commission. (2022). Guidance on a strategic framework for further supporting the deployment. https://environment.ec.europa.eu/topics/nature-and-biodiversity/green-infrastructure_en?prefLang=pt&utm_source=chatgpt.com#studies-and-publications Access 25 June 2026.

Forman, R. T. (2014). Urban ecology: science of cities. Cambridge University Press.

Fox, J. & Weisberg, S. (2019). An R Companion to Applied Regression, Third edition. Sage, Thousand Oaks CA.

Geospatial Information Authority of Japan. (2022). Basic Geospatial Information of water area. https://fgd.gsi.go.jp/download/menu.php. Access 29 May 2022

Gillies, CS., Beyer, HL., & St Clair, CC. (2011). Fine-scale movement decisions of tropical forest birds in a fragmented landscape. Ecological Applications, 21(3), 944–954. 10.1890/09-2090.1

Green, AJ., & Elmberg, J. (2014). Ecosystem services provided by waterbirds. Biological Reviews, 89, 105–122. 10.1111/brv.12045

Japan Aerospace Exploration Agency. (2025). ALOS Global Digital Surface Model “ALOS World 3D - 30m (AW3D30)“. https://www.eorc.jaxa.jp/ALOS/en/dataset/aw3d30/aw3d30_e.htm Access on 8 December.

Nagayama, S., & Nakamura, F. (2018). The significance of meandering channel to habitat diversity and fish assemblage: a case study in the Shibetsu River, northern Japan. Limnology, 19: 7–20. 10.1007/s10201-017-0512-4

Nakamura, F., Ishiyama, N., Sueyoshi, M., Negishi, J. N., & Akasaka, T. (2014). The significance of meander restoration for the hydrogeomorphology and recovery of wetland organisms in the Kushiro River, a lowland river in Japan. Restoration ecology, 22(4), 544–554. 10.1111/rec.12101

Nakano, D., & Nakamura, F. (2006). Responses of macroinvertebrate communities to river restoration in a channelized segment of the Shibetsu River, Northern Japan. River Research and Applications, 22(6), 681–689.10.1002/rra.928

Nakano, D., & Nakamura, F. (2008). The significance of meandering channel morphology on the diversity and abundance of macroinvertebrates in a lowland river in Japan. Aquatic Conservation: Marine and Freshwater Ecosystems, 18(5), 780–798. 10.1002/aqc.885

Phillips, R. A., Xavier, J. C., & Croxall, J. P. (2003). Effects of satellite transmitters on albatrosses and petrels. The Auk, 120(4), 1082–1090. 10.1093/auk/120.4.1082

R Core Team (2026). R: a language and environment for statistical computing. R Foundation for Statistical Computing. Vienna. https://www.R-project.org/. Accessed 1 September 2026

Rhoads, B. L., Schwartz, J. S., & Porter, S. (2003). Stream geomorphology, bank vegetation, and three dimensional habitat hydraulics for fish in midwestern agricultural streams. Water Resources Research, 39(8). 10.1029/2003WR002294

Shepard, E. L. (2025). How might turbulence affect animal flight in a changing world?. Journal of Experimental Biology, 228(Suppl_1), JEB248102. 10.1242/jeb.248102

Signer, J., Fieberg, J., & Avgar, T. (2019). Animal movement tools (amt): R package for managing tracking data and conducting habitat selection analyses. Ecology and evolution, 9(2), 880–890. 10.1002/ece3.4823

Skórka, P., Lenda, M., Martyka, R., & Tworek, S. (2009). The use of metapopulation and optimal foraging theories to predict movement and foraging decisions of mobile animals in heterogeneous landscapes. Landscape Ecology, 24(5), 599–609. 10.1007/s10980-009-9333-0

St. Clair, C. C., Bélisle, M., Desrochers, A., & Hannon, S. (1998). Winter responses of forest birds to habitat corridors and gaps. Conservation Ecology, 2(2). 10.5751/es-00068-020213

Takeshige, S., & Katoh, K. (2020). Usage of urban rivers by gulls and cormorants as movement pathways in winter. Ornithological Science, 19(2), 187–201. 10.2326/osj.19.187

Takeshige, S., & Katoh, K. (2023). Can rivers be important movement corridor for waterbirds in urban areas?. Landscape and Ecological Engineering, 19(3), 519–529. 10.1007/s11355-023-00557-7

Takeshige, S., Sawa, Y., & Katoh, K. (2025). Urbanization made rivers as important movement corridors for waterbird species, Black-headed Gulls. Landscape and Ecological Engineering, 21(3), 567–575. 10.1007/s11355-025-00658-5

Tremblay, M. A., & St. Clair, C. C. (2011). Permeability of a heterogeneous urban landscape to the movements of forest songbirds. Journal of Applied Ecology, 48(3), 679–688. 10.1111/j.1365-2664.2011.01978.x

Ward, J. (1998). Riverine landscapes: biodiversity patterns, disturbance regimes, and aquatic conservation. Biological conservation, 83(3), 269–278. 10.1016/S0006-3207(97)00083-9

Winemiller, K. O., Tarim, S., Shormann, D., & Cotner, J. B. (2000). Fish assemblage structure in relation to environmental variation among Brazos River oxbow lakes. Transactions of the American Fisheries Society, 129(2), 451–468. 10.1577/1548-8659(2000)129%3C0451:FASIRT%3E2.0.CO;2

