## Supplementary Information for "Structural characteristics of important daily movement corridors for waterbirds in urban areas: a field test focusing on Black-headed Gulls"

**The names of the authors**

**Title**

Structural characteristics of important daily movement corridors for waterbirds in urban areas: a field test focusing on Black-headed Gulls

**The affiliations and addresses of the authors**

1. Graduate School of Life Sciences, Tohoku University, 2-1-1 Katahira, Aoba, Sendai, Miyagi, 980-8577, JAPAN

2. Yamashina Institute for Ornithology, 115 Konoyama Abiko, Chiba, 270-1145, Japan

3. Faculty of Liberal Arts, The Open University of Japan, Wakaba 2-11, Mihama-ku, Chiba, 261-8586, Japan

**The e-mail address, and telephone number of the corresponding author**

Shiori Takeshige

+81-022-217-5054

**Supplementary Information**

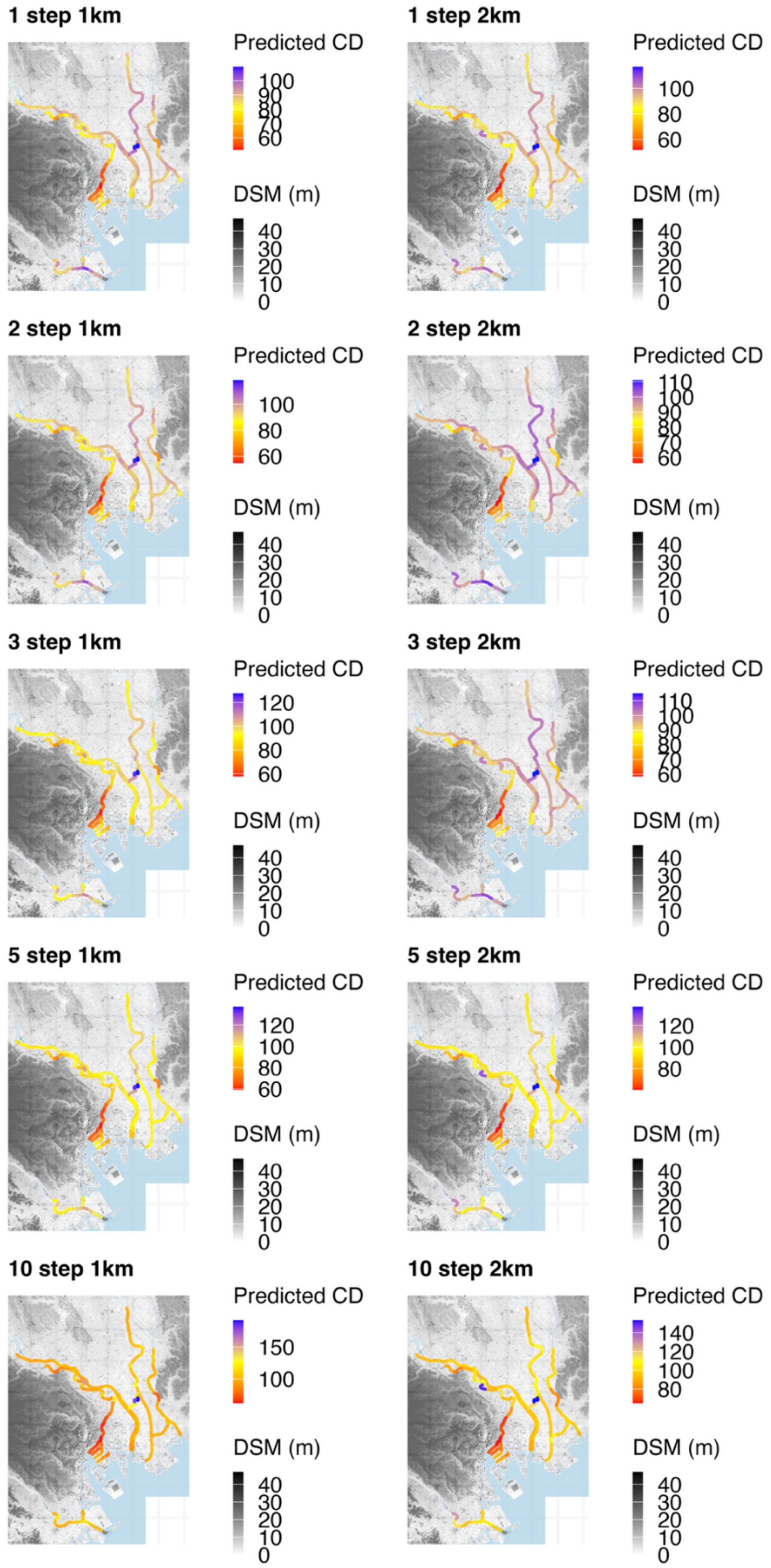

**Fig. S1** Predicted CD values at each river segment. Results are shown for the 1 and 2-km river segment scale. Warmer colors indicate a higher likelihood of gulls flying along the river segments. River segments where CD values could not be calculated due to excessive river width—which prevented the extraction of environmental variables—are shown in gray. Across all spatio-time scales, the results demonstrated that gulls consistently tended to fly along the river within the lower reaches of the Sumida River, which flows near the center of the map.
